# Iso-Seq enables discovery of novel isoform variants in human retina at single cell resolution

**DOI:** 10.1101/2024.08.08.607267

**Authors:** Luozixian Wang, Daniel Urrutia-Cabrera, Sandy Shen-Chi Hung, Alex W. Hewitt, Samuel W. Lukowski, Careen Foord, Peng-Yuan Wang, Hagen Tilgner, Raymond C.B. Wong

## Abstract

Recent single cell transcriptomic profiling of the human retina provided important insights into the genetic signals in heterogeneous retinal cell populations that enable vision. However, conventional single cell RNAseq with 3’ short-read sequencing is not suitable to identify isoform variants. Here we utilized Iso-Seq with full-length sequencing to profile the human retina at single cell resolution for isoform discovery. We generated a retina transcriptome dataset consisting of 25,302 nuclei from three donor retina, and detected 49,710 known transcripts and 241,949 novel transcripts across major retinal cell types. We surveyed the use of alternative promoters to drive transcript variant expression, and showed that 1-8% of genes utilized multiple promoters across major retinal cell types. Also, our results enabled gene expression profiling of novel transcript variants for inherited retinal disease (IRD) genes, and identified differential usage of exon splicing in major retinal cell types. Altogether, we generated a human retina transcriptome dataset at single cell resolution with full-length sequencing. Our study highlighted the potential of Iso-Seq to map the isoform diversity in the human retina, providing an expanded view of the complex transcriptomic landscape in the retina.

## Introduction

The human retina is a complex multi-layer tissue at the back of the eye, composed of a diverse range of specialized cell types with abundance varying from 0.05% to over 60% ^1^. Study of the genetic signals in different retinal cell types is important to understand the physiological processes that enable vision in humans. Recent advances in single cell RNAseq (scRNA-seq) offered an exciting opportunity to profile the transcriptomic landscape of different retinal cell types at a single cell level. Previous studies have generated a number of single cell transcriptomic datasets for human retina ^1–9^, as well as the ocular anterior segment ^10^ and posterior segment ^11^. Also, recent work from the Human Cell Atlas initiative has also compiled a large-scale human retina gene atlas with ∼2.4 million cells to improve the resolution of molecular profiling of the retina ^12^.

Current efforts of single cell transcriptome studies are limited in their ability to profile mRNA isoforms in a high-throughput manner, as the conventional 3’ sequencing only sequence ∼100bp base pairs which is unsuitable for mRNA isoform detection. mRNA isoforms can be derived from alternative processing of pre-mRNAs, including alternative splicing, use of alternative promoters, transcription start sites and/or polyadenylation sites. Over 92% of human genes undergo alternative splicing, representing a major mechanism that contributed to protein diversity and tissue-level regulation ^13^. Previous bulk RNA-seq studies revealed a high diversity of mRNA isoforms in human retina ^14,15^. However, comprehensive mapping of isoforms across major retinal cell types at a single cell level is currently lacking.

Recent advancement in the single-molecule real-time (SMRT) Iso-Seq technology by Pacific Biosciences (PacBio) enabled sequencing of full-length cDNA, providing an exciting opportunity to analyze the mRNA isoform landscape and map splice junctions. The PacBio long-read sequencing has been applied to bulk mouse retina to identify isoform and long noncoding RNA expression, yielding a dataset of ∼372k circular consensus sequences (CCS) ^16^. Also, Ray et al. has applied targeted long-read sequencing in the bulk human retina to profile isoforms of the retinal degeneration gene *CRB1* ^17^. In this study, we utilized IsoSeq for isoform discovery in the human retina at a single cell level. Our retina transcriptome dataset consisted of 25,302 nuclei in major retinal cell types, with ∼1.4 billion short-reads and >24 million CCS reads, representing 59x more full-length sequencing depth compared to the previous study of bulk long-read sequencing in mouse retina ^16^.

## Results

### Isoform discovery in human retinal cell types

We collected three retina/choroid samples from three healthy donors (Supplementary table 1). Using the Chromium system (10X Genomics), we captured 25,302 nuclei from 3 donor sample with ∼8,000 nuclei/sample. The full-length cDNA library was then divided and processed for short-read sequencing using Novaseq following the standard 10X Genomics 3’ sequencing protocol, as well as long-read sequencing using Pacbio Sequel II. In total, we generated ∼1.4 billion short-reads (>55,000 reads/cell) and >24.1 million HiFi reads with >99% accuracy using 9 SMRT 8M cells (Figure 1A). Using the short-reads, we annotated 10 major cell types representative of the retina and the choroid/sclera layers, including rods, cones, Muller glia, astrocytes, horizontal cells, bipolar cells, amacrine cells, RPE, smooth muscle cells and fibroblasts (Figure 1B, 1C). We did not detect any obvious bias for donor cell distribution (Supplementary figure 1A) in the dataset, and most nuclei were covered by both short-read and long-read sequencing (Supplementary figure 1B). Following mapping, our long-read dataset contained ∼22M CCS reads and identified 34,936 unique genes (Supplementary table 1). We detected 291,721 unique isoforms, including 49,710 known isoforms and 241,949 novel isoforms. Altogether 243,236 splice junctions were detected, with 160,173 as known canonical splice junctions and 83,031 as novel canonical junctions.

**Figure 1:**
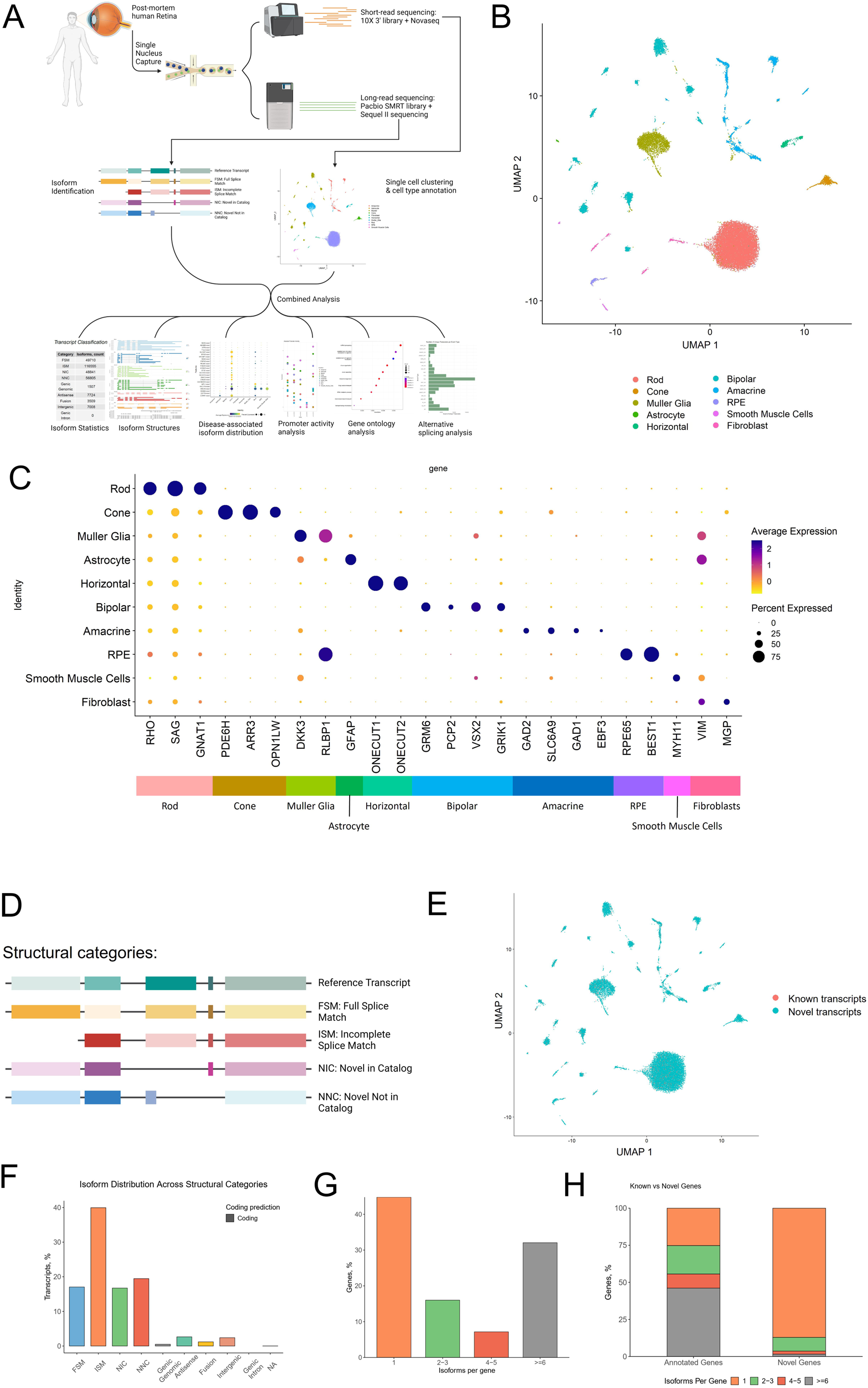
Isoform discovery in human retina at single cell resolution. A) Schematic of Iso-Seq of human retina and choroid at single cell resolution. B) UMAP plot of major retinal cell types. C) Dotplot showing expression of marker genes in retinal cells. D) Schematic of the structural categories of long-read transcripts based on SQANTI classification. E) UMAP plot showing the distribution of known isoforms (FSM) and novel isoforms (ISM+NIC+NNC) in retinal cells. F) Isoform distribution across structural categories. Summary of G) isoforms per genes, and H) novel and known gene proportions in the human retina long-read dataset.

Structural characterization of isoforms was performed using the *pigeon* pipeline (https://isoseq.how/classification/pigeon.html). According to SQANTI classification (Figure 1D), transcripts mapping a known reference (FSM) accounted for 17.0% of the sequencing data, while novel transcripts of known genes (ISM, NIC, NNC) made up 76.2% of the dataset (Figure 1F). Other transcripts consist of those that are ‘genic-genomic’ (partial exon and intron/intergenic overlap in a known gene), antisense (overlapping the complementary strand of known gene, fusion (spanning two annotated loci) or intergenic (transcript outside boundaries of annotated gene) (Supplementary figure 2). The median transcript lengths for FSM, ISM, NIC and NNC are 1.48kb, 1.38kb, 1.64kb, and 1.49kb respectively (Supplementary figure 2C-E). On average we observed higher exon numbers in the novel isoforms (NIC, NNC and ISM) compared to the known isoforms (FSM) (Supplementary figure 2F-H). Among the splice junctions detected, 5.8% and 26.6% were novel canonical splice junctions in NIC and NNC respectively, and we did not detect any novel splice junctions in FSM and ISM as expected (Supplementary figure 3A).

Next, we integrated orthogonal genomic annotations to the long-read dataset. At the 5’ end, the transcription start site is detected within ±100bp and ±500bp of the annotated site for FSM and ISM respectively (Supplementary figure 3B-C). Furthermore 55% of novel isoforms were supported by proximal annotated CAGE (Supplementary figure 4A-B). At the 3’ end, 39% of novel isoforms were supported by polyA motifs (Supplementary figure 4C). For ISM, most of the polyA motifs were detected >5kb from the annotated site, which is indicative of the presence of novel isoforms (supplementary figure 3E). PolyA distance analysis showed that the polyA motif was most commonly detected at ∼15bp from the 3’ end in FSM, ISM, NIC and NNC (Supplementary figure 4D).

Notably, 32.1% of detected genes have >6 isoforms and they are mostly annotated genes, highlighting the abundance of isoform variants in the retina (Figure 1G-H). On the other hand, 7.2% of detected genes have 4-5 isoforms and 16.0% of genes have 2-3 isoforms (Figure 1G). Gene ontology analysis showed that the top GO processes for known isoforms (FSM) and novel isoforms (ISM+NIC+NNC) were ncRNA processing, protein localization and cilium assembly (Supplementary figure 1C-F). In addition, GO processes for known isoforms also included macroautophagy and mitochondrial gene expression, whereas GO processes for novel isoforms included Golgi vesicle transport and microtubule-based transport.

### Differential use of alternative promoters to drive isoform expression in retinal cells

Next we studied the use of alternative promoters to drive isoform expression. Our results showed that ∼1-8% of genes utilized multiple promoters across major retinal cell types. Bipolar cells have the highest percentage of genes with multiple promoters (8.3%), followed by amacrine cells (7.8%) and Muller glia (7%). We also analyzed the relationship between promoter activity and their position. In general, we observed that major promoters are more frequently featured with decreasing promoter position, and vice versa for minor promoters (Figure 2B). This phenomenon is more prominent in amacrine cells, cones and rods, but less prominent in astrocytes and smooth muscle cells (Figure 2B). For instance, *AHI1*, an IRD gene involved in recessive Joubert syndrome ^18^, is predominantly expressed using pmtr.32032 as the major promoter, while ptmr.32025 represents a minor promoter with lower activity (Figure 2C). We observed this preferential use of alternative promoter in bipolar cells, cones, Muller glia cells, rods, and smooth muscle cells.

**Figure 2:**
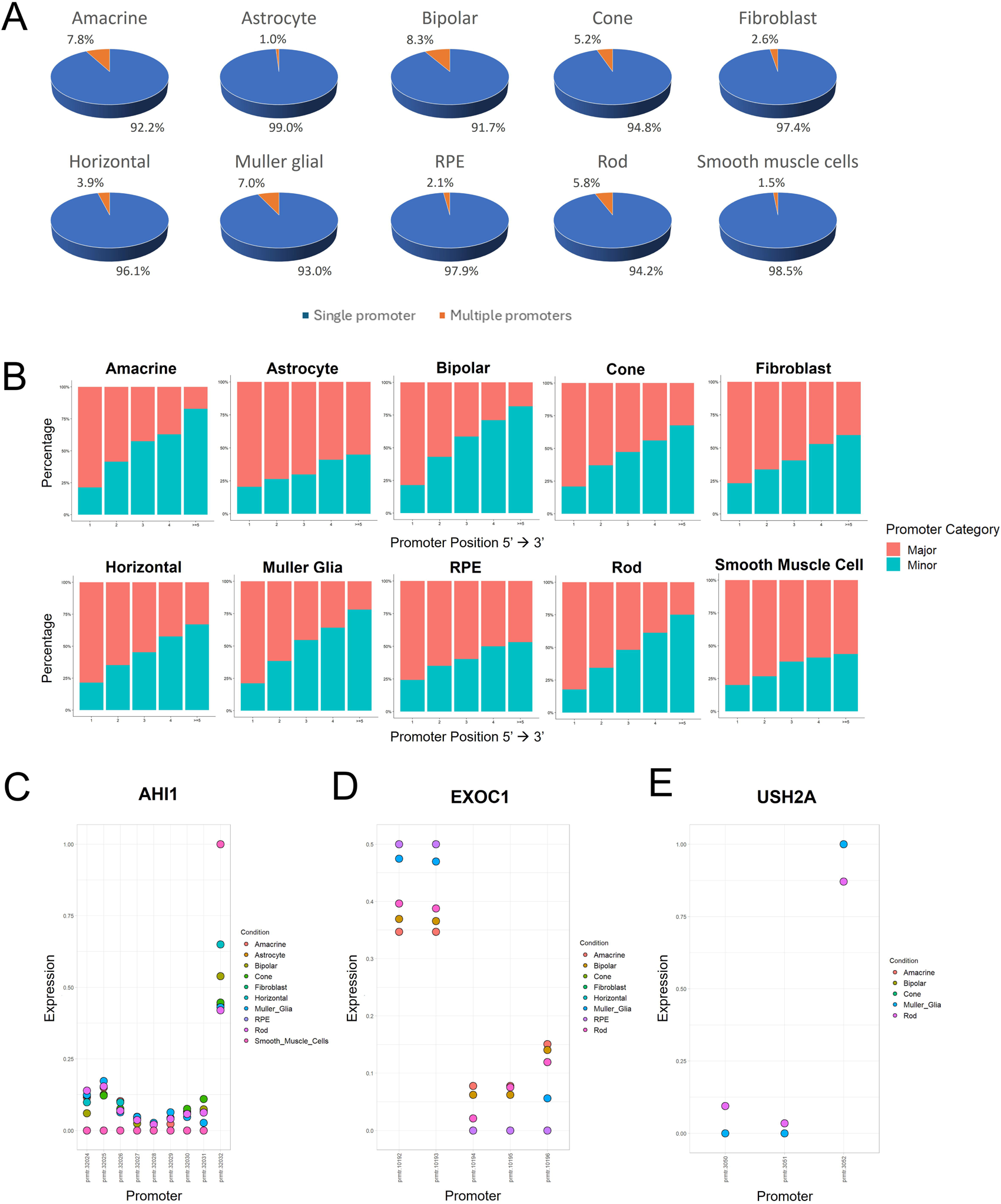
Differential use of alternative promoters in retinal cells. A) Pie chart showing the proportion of genes using single or multiple promoters in retinal cells. B) Analysis of promoter activity and their position in retinal cells. Proportions of major promoters are depicted in blue, proportion of minor promoters are depicted in red. C-E) Relative promoter activities for C) *AHI*, *EXOC1*, E) *USH2A* in major retinal cell types.

Also, our results highlighted the differential use of alternative promoters to drive gene expression in different retinal cell types. We compared the major promoter activity and gene expression and showed that a single major promoter does not fully explain gene expression in retinal cell types, with minor promoters also contributing to gene expression (Supplementary figure 5). For example, *EXOC1* expression is solely driven by pmtr.10192 and pmtr.10193 in RPE, while its expression can also be driven by pmtr.10196 in amacrine cells, bipolar cells, Muller glia cells and rods (Figure 2D). Similarly, *USH2A* expression is driven by a single promoter in Muller glia (ptmr.3052), while its expression can be driven by multiple promoters in rods (Figure 2E).

### Identification of novel isoforms for genes responsible for inherited retinal diseases

Using our long-read dataset for isoform discovery, we analyzed the expression pattern of novel isoforms for genes associated with various forms of retinal degeneration from RetNet, including retinitis pigmentosa (RP), cone-rod dystrophy (CORD), macular degeneration, congenital stationary night blindness (CSNB), Leber congenital amaurosis (LCA) and Usher Syndrome.

CORD is a group of inherited retinal diseases characterized by progressive degeneration of the cones followed by the rods in the retina, leading to severe vision impairment ^19^. The inheritance pattern can be autosomal dominant, autosomal recessive or X-linked. Notably, in the cones we detected novel isoforms for a number of CORD genes (e.g. *GNAT2, GUCA1A*), while some CORD genes were expressed in both cones and rods (e.g. *UNC119, AIPL1;* Figure 3A).

**Figure 3:**
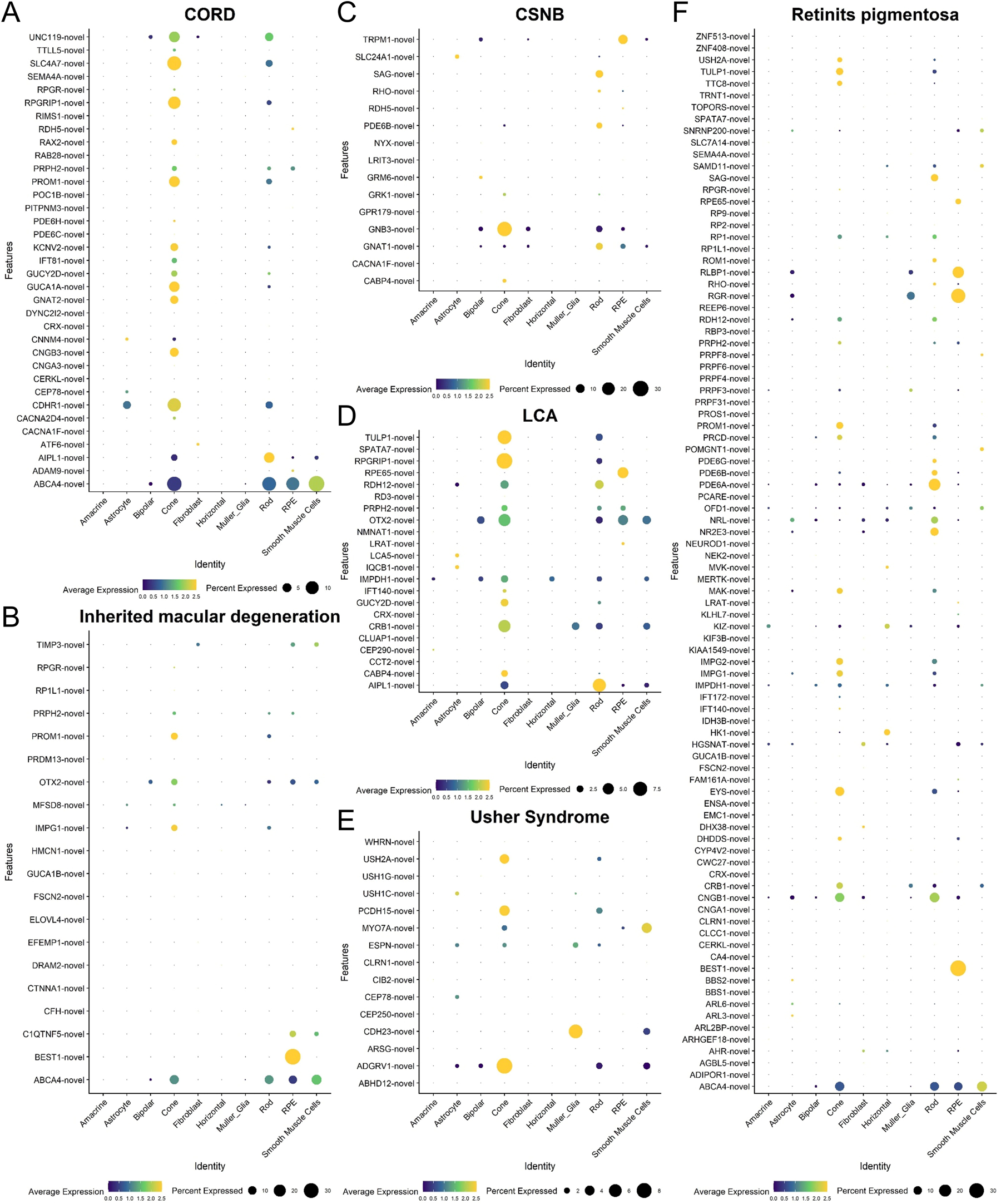
Identification of novel isoforms for IRD genes in retinal cell types. Dot plots showing novel isoform detection of genes associated with A) cone-rod dystrophy (CORD), B) inherited macular degeneration, C) congenital stationary night blindness (CSNB), D) Leber congenital amaurosis (LCA), E) Usher Syndrome and F) retinitis pigmentosa (RP).

Macular degeneration such as Stargardt disease, vitelliform degeneration, Sorsby macular dystrophy and familial drusen, causes gradual retinal degeneration leading to central vision loss in the patients ^19^. Our results identified novel isoforms of macular degeneration genes in photoreceptors (eg. *PROM1, IMPG1*; Figure 3B). In addition, in RPE we detected novel isoforms of the Best disease gene *BEST1*, as well as *C1QTNF5* which causes Late-onset retinal degeneration (L-ORD). We also discovered abundant novel isoforms of the Stargardt disease gene *ABCA4* in RPE and photoreceptors.

CSNB is an IRD that affects vision in low-light conditions from birth. Unlike other forms of night blindness, CSNB is non-progressive and typically remains stable ^20^. Our data detected novel isoforms of CSNB genes in rods (eg. *SAG, GNAT1;* Figure 3C), as well as *GNB3* in both rods and cones which potentially play a role in pathogenesis of CSNB.

LCA is a rare IRD that causes photoreceptor dysfunction, resulting in severe visual impairment from birth or early age ^19^. LCA affects both the rod and cone function. Consistent with this, we detected a number novel isoforms of LCA genes in cones and rods (eg. *TULP1, RPGRIP1*, *CRB1;* Figure 3D).

Usher syndrome is characterized by a combination of hearing loss and visual impairment, with varying degrees of sensory loss ^21^. Our results highlighted the presence of novel isoforms of *USH2A* and *ADGRV1* in cones, as well as *CDH23* in Muller glia (Figure 3E).

Finally, a number of genes have been identified to cause RP - a disease characterized by degeneration of both the rods and cones ^22^. We observed the presence of a number of novel RP gene variants in rods (eg. *PDE6A, PDE6B, PDE6G;* Figure 3F), cones (eg. *TTC8, TULP1, EYS*), or both cones and rods (eg. *CNGB1*). Altogether, these results suggest an understudied role of novel isoforms in contributing to IRD pathology.

### Isoform profiling and differential exon splicing in retinal cells

To demonstrate the potential of our dataset to profile isoforms in the human retina, we analyzed the isoform distribution of a IRD gene *ABCA4* that causes Stargardt disease, which is a highly polymorphic gene with over 1200 disease-associated variants ^23^. Of the likely pathogenic variants, 18% of them were determined to have a significant splice site alteration ^24,25^. In the retina/choroid sample, we detected high *ABCA4* expression in rod, cones, RPE and smooth muscle cells, and to a lesser extent in Muller glia and bipolar cells (Figure 4A). By analyzing full-length transcripts with CAGE peaks and poly-A tails, we identified many *ABCA4* full-length isoforms (Figure 4B). The rods showed the highest number of *ABCA4* isoforms, followed by cones, bipolar cells, MG cells, smooth muscle cells and RPE cells, whereas single isoform is detected in amacrine cells and horizontal cells. Also, we detected a range of novel exons for *ABCA4* variants, with >40 novel exons in cones, rods and RPE (Supplementary figure 6). We selected a representative isoform expressed in rods, cones, RPE and smooth muscle cells and predicted the protein structures using AlphaFold 3 ^26^. Our results highlighted the diverse variations in protein structure exhibited in these *ABCA4* isoforms (Figure 4C), which was contributed by the use of novel splicing junctions in the *ABCA4* gene (Figure 4D). Together, these results highlighted the presence of a diverse profile of *ABCA4* variants in retinal cell types.

**Figure 4:**
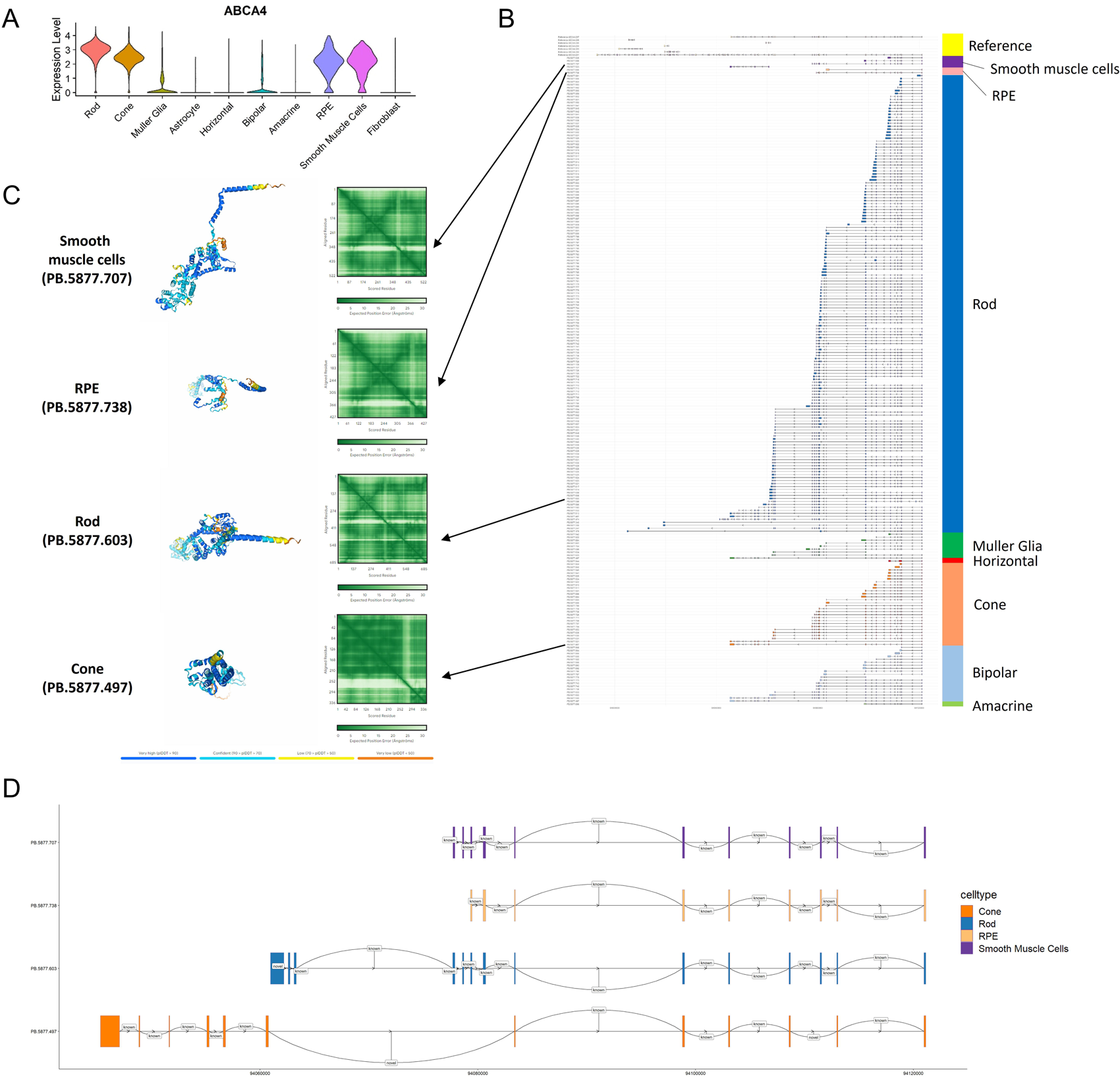
Transcript diversity of the IRD gene *ABCA4* in human retinal cells. A) Violin plot of *ABCA4* expression in major retinal cell types using the short-read sequencing data for quantification. B) Profiling of *ABCA4* full length isoforms with CAGE peaks and poly-A tails in in smooth muscle cells (purple), RPE (pink), rods (dark blue), Muller glia (dark green), horizontal cells (red), cones (orange), bipolar cells (light blue), amacrine cells (light green) compared to annotated isoforms (yellow). C) AlphaFold 3-predicted protein structure of representative ABCA4 isoforms in smooth muscle cells, RPE, rods and cones, and D) identified novel and known splicing junctions in the isoforms.

Next, we categorized the alternative splicing events based on ASprofile classification ^27^. We showed that the most common alternative splicing events in the retina utilized alternative transcription start sites (TSS), followed by exon skipping (SKIP) and alternative exon ends (AE, Figure 5A). Also, we analyzed the differential exon usage in retinal cells and measured the relative abundances of the splicing events as the percentage spliced-in (PSI). Our results identified 131 significant exons that were differentially spliced by specific retinal cell types (Figure 5B). The top genes that were differentially spliced include *PNISR*, *VDAC3* and *AHI1* (Supplementary data 1). Also, we categorized the retinal cells into cell groups and assess differential exon usage. We identified 31 significant differentially spliced exons between neurons and non-neurons, 30 between glial cells and non-glial cells, and 14 between photoreceptors and other neurons (Figure 5C). The top significant spliced exons are mapped to *VDAC3*, *ZNF536* and *ASPH* (Supplementary data 1). For example, *ASPH* is implicated in Traboulsi syndrome which is characterized by facial dysmorphia and complex ocular abnormalities ^28^. We detected 68 isoforms of *ASPH* across the retinal cell types, with high isoform diversity observed in the rods, MGs and the amacrine cells (Supplementary figure 7). *ASPH* has >30 annotated exons and our results showed preferential usage for an annotated exon in amacrine cells, horizontal cells and rods (all PSI>0.95) compared to Muller glial cells (PSI=0.052; Figure 5D). In another example, *EXOC1* encodes for exocysts that are involved in photoreceptor ciliogenesis ^29^. Using our long-read dataset, we identified 30 isoforms of *EXOC1* across the retinal cell types (Supplementary figure 8). In particular, our results highlighted a novel exon of *EXOC1* that is preferentially used in amacrine and rod (both PSI=1) compared to Muller glial cells (PSI=0.037; Figure 5E), highlighting an expanded transcriptome complexity in the retina.

**Figure 5:**
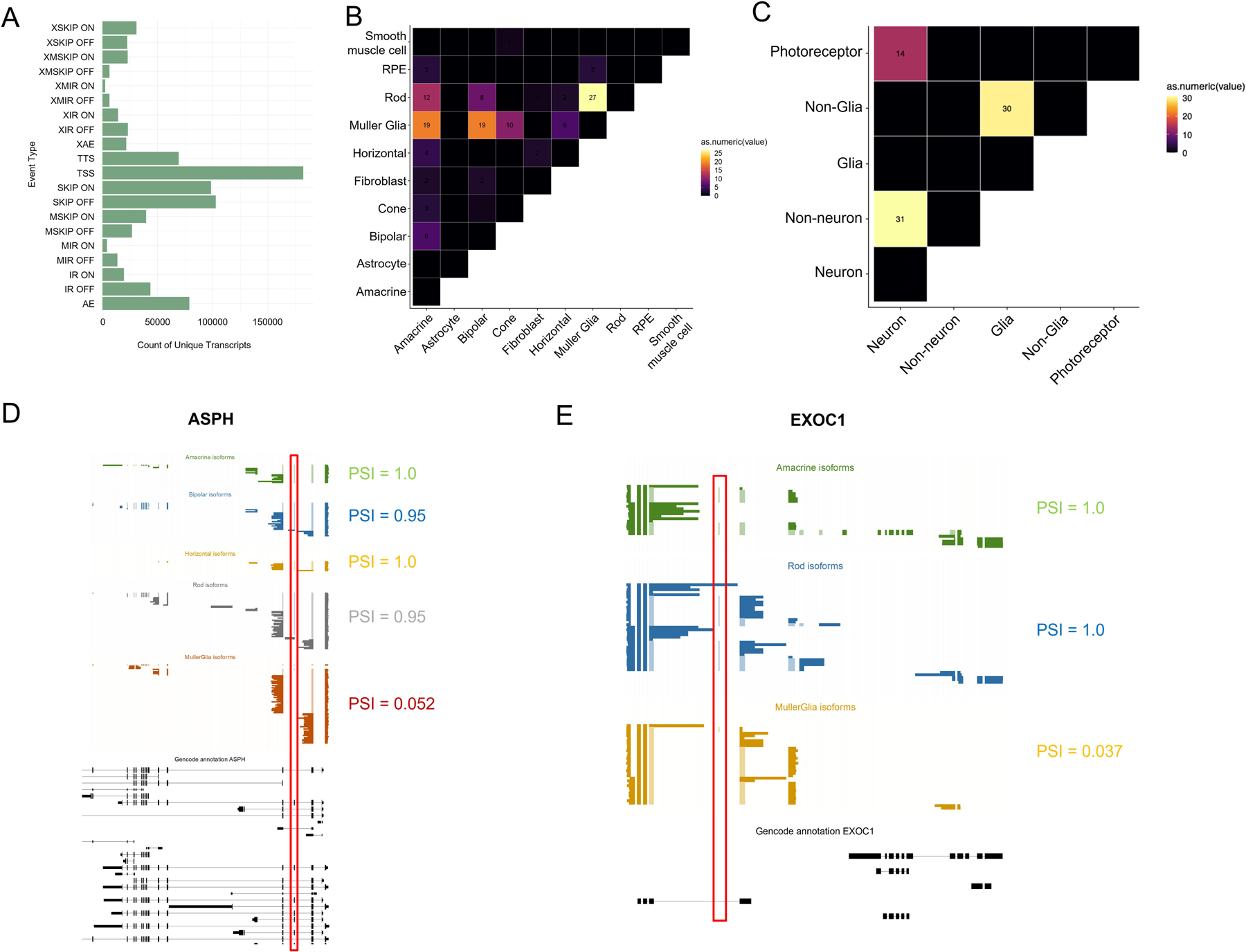
Alternative splicing in retinal cell types. A) Categorization of alternative splicing events in the retina, including SKIP (exon skipping), MSKIP (cassette exons), IR (retention of single introns), MIR (retention of multiple introns), AE (alternative exon ends), TSS (alternative transcription start site), and TTS (alternative transcription termination site). B) Heatmap illustrating the number of significant exons that were differentially spliced in retinal cell types. Black squares indicated cell type comparison that were not tested. C) Heatmap illustrating the number of significant exons that were differentially spliced in retinal cell groups (neurons, non-neurons, glial cells, non-glial cells, photoreceptors). Black squares indicated cell type comparison that were not tested. D) Representative example of an annotated exon in *ASPH* that was differentially spliced in Muller glia (brown) compared to other retinal cell types: Amacrine cells (green), bipolar cells (blue), horizontal cells (gold), rods (grey). Annotated isoforms of *ASPH* were displayed in black. Red rectangle marked the differential spliced exon. Representative example of a novel exon in *EXOC1* that was differentially spliced in Muller glia (gold) compared to amacrine cells (green) and rods (blue). Annotated isoforms of *EXOC1* were displayed in black. Red rectangle marked the differential spliced exon.

## Discussion

Conventional single cell transcriptomics based on short-read sequencing has been widely used to survey the transcriptomic landscape in healthy or diseased human retina. Recent advances in Iso-Seq offers complete characterization of full-length transcripts, representing a powerful tool to identify transcriptional start and end sites, alternative splicing and splice junctions. Iso-Seq has been used for isoform profiling from a range of human tissues, including cerebral cortex, lymphocytes and gastric cancer ^30–35^. Here we leveraged Iso-Seq using PacBio long-read sequencing to comprehensively map isoform expression in human retina at a single cell level. Our results identified 241,949 previously unannotated isoform transcripts across major retinal cell types, highlighting the complex transcript diversity in human retina.

Many genes can be transcribed into distinct isoform variants using different alternative promoters, exon arrangements, and 3’ untranslated regions. Understanding of the transcript element combinations is important to fully understand gene regulation in the retinal cell types. Our results showed that up to 8% of genes utilized alternative promoters across major retinal cell types, with the highest activity observed in bipolar cells. For example, *AHI1* is implicated in Joubert Syndrome characterized by neurological and retinal abnormalities ^18^. A knockout study in mice showed that *Ahi1* is critical for photoreceptor development and plays a role in stabilizing the outer segment proteins ^36^. Our results showed the use of alternative promoters in driving isoform diversity for *AHI1* in rods, cones, MG, bipolar and amacrine cells, highlighting an understudied role of *AHI1* isoforms in the retinal neurons. Furthermore, we confirmed the importance of alternative splicing in contribution to transcript diversity in the retina. We showed differential exon usage in major retinal cell types and identified 161 significant exons that were differentially spliced in retinal genes, such as *ASPH* and *EXOC1*. Notably, we identified many novel isoforms of known IRD genes across different retinal cell types, providing novel insights into the transcriptomic complexity in the human retina. For example, we identified 166 isoforms for *ABCA4*, an IRD gene that caused Stargardt disease. *ABCA4* is a highly polymorphic gene with 50 exons and >1200 disease associated variants ^23^. Fujinami et al. previously reported that 18% of the pathogenic variants contain splicing site alteration ^24^. Our results expanded this view and highlighted the transcript diversity of *ABCA4* in different retinal cell types. It is possible that some of these novel isoforms for IRD genes contribute to disease pathogenesis. Functional studies of the identified novel isoforms in the future would be important and could identify novel drug targets to develop treatment for the corresponding IRD.

There are some limitations in our study. Firstly, our dataset excluded the retinal ganglion cells, possibly due to the low number of ganglion cells present in our subsampling of retina tissue. To address this, future studies could utilize cell enrichment using specific markers to better capture rare cells such as retinal ganglion cells, a strategy that has yielded some success in previous scRNA-seq studies ^37^. Secondly, the long-read sequencing depth in this study is suitable for isoform discovery but is limited for comprehensive quantification of isoform levels. Although the long-read sequencing depth in our dataset represented a significant improvement to previous report in the mouse retina ^16^ (59x more sequencing depth), it is likely that higher sequencing depth would improve the statistical power to identify more differential splicing, exon usage and isoform expression in the retinal cells. Recent advances in long-read sequencing technology to improve the throughput and lower the cost would help achieve this, such as the development of concatenation protocol to improve reads output for CCS for PacBio sequencing ^38,39^. Also, future ISO-seq studies to profile retina from donors with different ages and ethnicity would be important to expand on the sample diversity.

In summary, here we report a cell-specific atlas of isoform expression for human retina, providing a useful resource to the scientific community for retinal biology study. Our work provides an expanded view on the complex transcriptomic landscape in the human retina, and highlights the potential of using long-read sequencing for isoform discovery in highly heterogeneous tissue.

## Methods

### Donor sample collection

Donor sample collection was approved by the Human Research Ethics committee of the Royal Victorian Eye and Ear Hospital (HREC13/1151H and HREC22/1555H) and carried out in accordance with the approved guidelines. Donor consent was obtained and the experiments conformed to the principles set out in the WMA Declaration of Helsinki. Post-mortem eye globes were collected by the Lions Eye Donation Service (Royal Victorian Eye and Ear Hospital, Melbourne, Australia) and the retina/choroid was dissected for this study. Following dissection, the sample was snap-frozen in isopentane and liquid nitrogen for storage.

### Single-nuclei isolation and library construction

Retinal tissue was dissociated and single nuclei isolation was performed using snRNA-seq protocol described previously ^40^. Single nuclei from three retina samples were stained with DAPI and sorted using a FACS Aria II, followed by processing with the Chromium system for single nuclei capture (10X Genomics). For short-read sequencing, barcoded cDNA libraries were prepared using the single-cell 3’ mRNA kit (V3.1; 10X Genomics). The quality of the cDNA libraries were checked using TapeStation and processed for 150bp paired-end sequencing using Novaseq (Australian Genome Research Facility).

For long-read sequencing, the Pacbio protocol for single cell ISO-seq libraries using SMRTbell Express Template Prep Kit 2.0 was performed (Australian Genome Research Facility). The barcoded cDNA libraries were re-amplified for 6 cycles to provide enough cDNA for downstream Pacbio library preparation. Library preps were QC using Bioanalyzer and processed for Sequel II sequencing using 9 SMRT 8M cells.

### Data processing and quality control of transcriptomic data

For short-read data, data processing and quality control were performed using the standard 10x Genomics cellranger pipeline (v6.0.1) as previously described ^2,9^. Briefly, fastq files were generated from the raw Illumina BCL files using *mkfastq*. *Cellranger count* was used to generate read count matrices and mapped to reference genome (GRCh38). The count data were imported to *Seurat* (v3) for downstream analysis ^41^. Quality control was performed to retina high-quality outlier nuclei with <5% mitochondria content, >200 genes and <5000-6000 genes (sample dependent). Genes were retained in the data if they were expressed in >3 nuclei.

For long-read data, the Iso-seq3 (v4.0.0) pipeline was used to process the raw sequencing runs to generate deduplicated reads for transcript classification and tertiary analysis ^42^. We performed single-molecule circular consensus sequencing (CCS) to generate HiFi reads which provide accuracy of 99.9% from productive zero-mode waveguides (ZMWs). The removal of primers and identification of barcodes of HiFi reads was performed using *lima* to generate full-length (FL) reads. Then the tags such as UMIs and cell barcodes, as well as full-length tagged (FLNC) reads which were refined by trimming of polyA tails and unintended concatemer identification and removal, were clipped and refined by *isoseq tag* and *isoseq refine* functions using default settings, with the *--require-polya* parameter to filter for FLNC reads that have a polyA tail with at least 20 base pairs. The FLNC reads from 9 SMRT cells were merged together and the cell barcode whitelist of 10x single-cell 3’ libraries was then used to reassign erroneous barcodes based on edit distance to correct identified cell barcode errors and estimates which reads were likely to originate from real cells. Lastly, PCR deduplication via clustering by UMI and cell barcodes was performed by the *isoseq groupdedup* function.

### Transcript classification of FLNC reads

Deduplicated FLNC reads were mapped to the reference genome (GRCh38) using *pbmm2* which is a minimap2 frontend for PacBio native data formats. The mapped reads were then collapsed into unique isoforms based on exonic structures using *isoseq collapse*. These isoforms were classified into different structural categories using *pigeon* (v1.0.0). Supplemental reference information of the identified isoforms, including CAGE peaks, intropolis junctions, polyA sequences, and FLNC counts, were added to the classification output following the *pigeon classify* function. The *pigeon filter* function was performed to filter isoforms from the classification output and the *pigeon make-seurat* function was used to construct isoform count matrix for downstream Seurat process. The filtered classification results were summarized in a pigeon report using the ‘*pigeon_report.R*’ script distributed by the SQANTI3 package ^43^. Full splice match (FSM) molecules are considered known transcripts, and other transcripts are considered novel transcripts (ISM, NIC and NNC).

### Dimension reduction, clustering, and cell-type annotation

The isoform count matrix of PacBio long-read data generated by *pigeon* and the gene count matrix of single-cell short-read data generated by *cellranger* were imported into Seurat (v3) for tertiary analysis ^41^. The short-read data for individual retina samples was normalized using log transformation and variance stabilization, and the three samples were merged based on integration anchors defined in Seurat. Linear dimensionality reduction using principal components analysis (PCA) was performed, and a K-nearest neighbor (KNN) graph was generated based on the Euclidean distance in the PCA space to calculate the neighborhood overlap between each cell and its local neighbors. The Louvain algorithm was used for modularity optimization via *FindClusters* to iteratively group cells together at a resolution of 0.1 and 0.5. Uniform manifold approximation and projection (UMAP) non-linear dimensional reduction algorithm was applied in Seurat to place similar cells together in the low-dimensional space. Cell types were annotated using known canonical marker genes for retinal cell types as previously described^2,9^. For long-reads, genes detected with at least two counts were retained in the isoform count matrix and pseudo-bulk to determine the isoform expression according to the cell type. The isoform count matrix was embedded to the short-read Seurat object as an ‘isoform’ assay using barcode deconvolution. Meanwhile, the transcript classification and annotation generated previously was added to the metadata slot of the combined Seurat object to help categorize the isoforms into different structure classes. The cell type of each cell in the long-read and short-read combined Seurat object was transferred from the short-read analysis through barcode matching.

### Alternative promoter analysis

The isoform count matrix was pseudo-bulked and subsetted according to their cell identities. The barcode list of single cells and their cell types was used to subset the deduplicated and mapped *BAM* files. The cell type-specific long-read sequencing transcriptomes were input into proActiv (v0.1.0) together with the promoter annotation file (GRCh38 Gencode v44) to identify alternative promoter usage ^44^. Identified promoters were categorized into different classes: 0.25 was used as the threshold to identify active and inactive promoters, and the most active promoters of each transcript were classified as major promoters while the rest with a lower level activity were categorized as minor promoters.

### Alternative splicing analysis

The transcript *GFF* file was collapsed, sorted, and filtered using the *isoseq collapse*, *pigeon prepare*, and *pigeon filter --isoforms* function following the Iso-seq3 (v4.0.0) pipeline. The alternative splicing events were identified and classified using ASprofile (v1.0.4) ^27^. The alternative splicing event class described in this study includes SKIP (exon skipping), MSKIP (cassette exons), IR (retention of single introns), MIR (retention of multiple introns), AE (alternative exon ends), TSS (alternative transcription start site), and TTS (alternative transcription termination site). The output file was loaded into R for statistical analysis and further visualization.

### Identification and visualization of specific isoforms

The gene lists associated with retinal degenerative diseases were collected from the Retinal Information Network (RetNet) ^45^. Information such as the count matrix, cell types, transcript structures, isoform classifications, alternative splicing events, and gene ontologies were imported into Seurat for further visualization. Also, the *GFF* file used in the alternative splicing event classification was filtered for transcripts with CAGE sites and polyA tails, and then loaded into *ggtranscript* (v0.99.9) to visualize the detailed gene structure of transcript variants ^46^.

### Differential exon usage

Sequenced reads were aligned using STAR^47^ on genome hg38 and barcode information was taken from the short-read sequencing dataset. The scisorseqr (v0.1.9, ^48^) package was used to pre-process the data and the *casesVcontrols* function from scisorATAC^49^ package was used with standard settings to identify alternative exons.

### Protein structure prediction

The full length sequence of the isoform was extracted and input into NCBI ORFfinder (https://www.ncbi.nlm.nih.gov/orffinder) to identify the ORF, using a minimal ORF length of 75 nucleotides and start codon as ATG and alternative initiation codons. Protein structure prediction was performed using AlphaFold 3 ^26^, using the longest ORF as input with default setting.

### Data availability

The code for data analysis is available at Github (https://github.com/wongcb/Human-retina-ISOseq). The sequencing data in this study are available in the Human Cell Atlas data portal.

## Author contribution

RCBW and HT designed the experiments; DUC, SH and RCBW conducted the experiments; LW, SL, CF, HT and RCBW analyzed the data; RCBW, AH, PYW and HT provided funding to this work; LW and RCBW wrote the manuscript. All authors approved the manuscript.

## Acknowledgement

We thank Dr Casey Anttila and the SCORE facility (Walter and Eliza Hall Institute of Medical Research) for technical assistance with flow cytometry and 10X library preparation. RCBW is supported by the University of Melbourne, the Centre for Eye Research Australia, National Health and Medical Research Council (GCT1184076) and Medical Future Research Fund (MRF2024365). The Center for Eye Research Australia receives Operational Infrastructure Support from the Victorian State Government.

