## Supplementary Info: Suppl figure 1-8, suppl table 1 for "Iso-Seq enables discovery of novel isoform variants in human retina at single cell resolution"

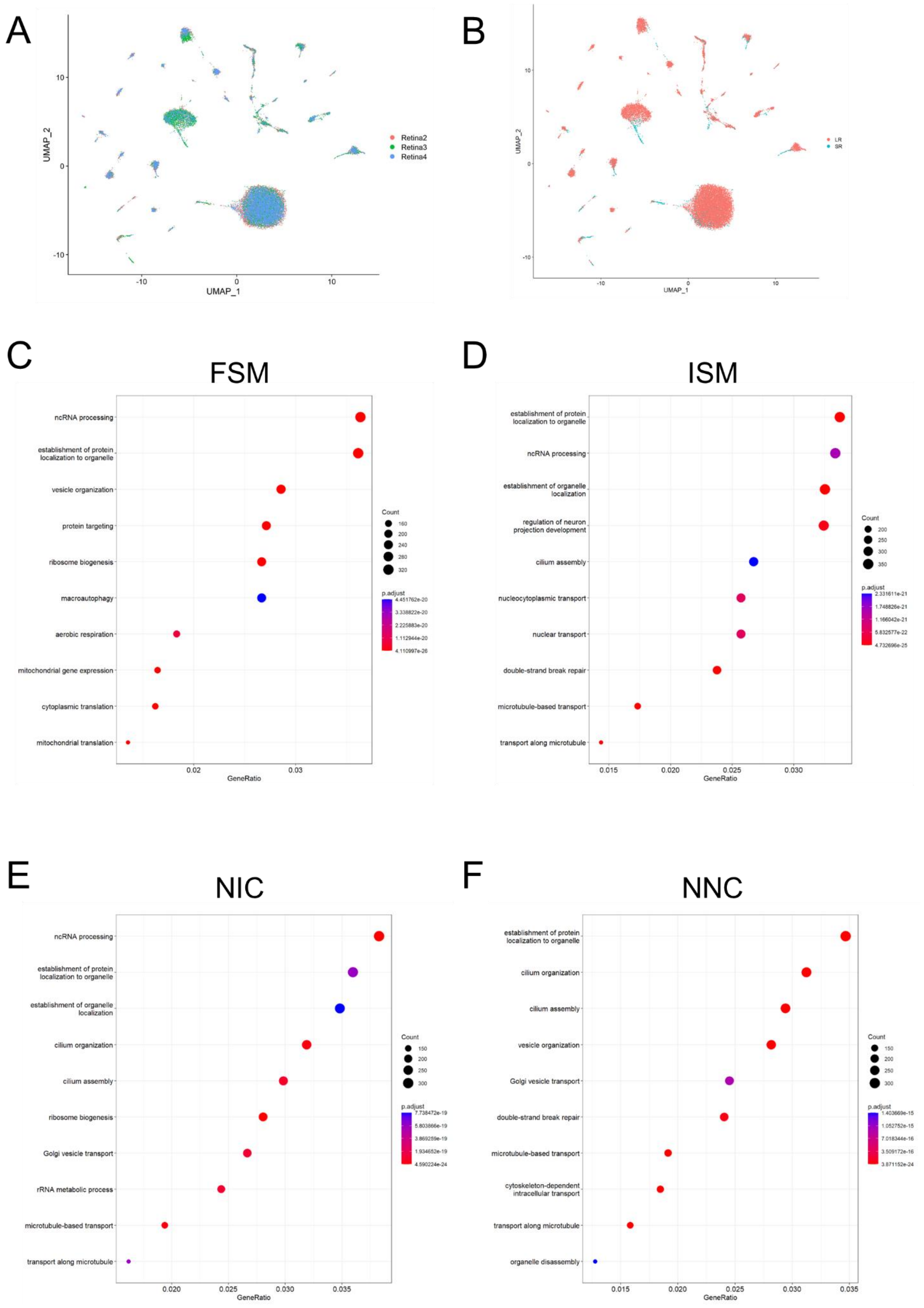


**Supplementary figure 1: Characterization of the retina Iso-Seq dataset.** Feature plot showing the sample distribution of A) donor retina (Retina 1-3), and B) coverage for short-reads (SR) and long-reads (LR). C-F) Gene ontology analysis of C) FSM, D) ISM, E) NIC and F) NNC identified in the long-read dataset.

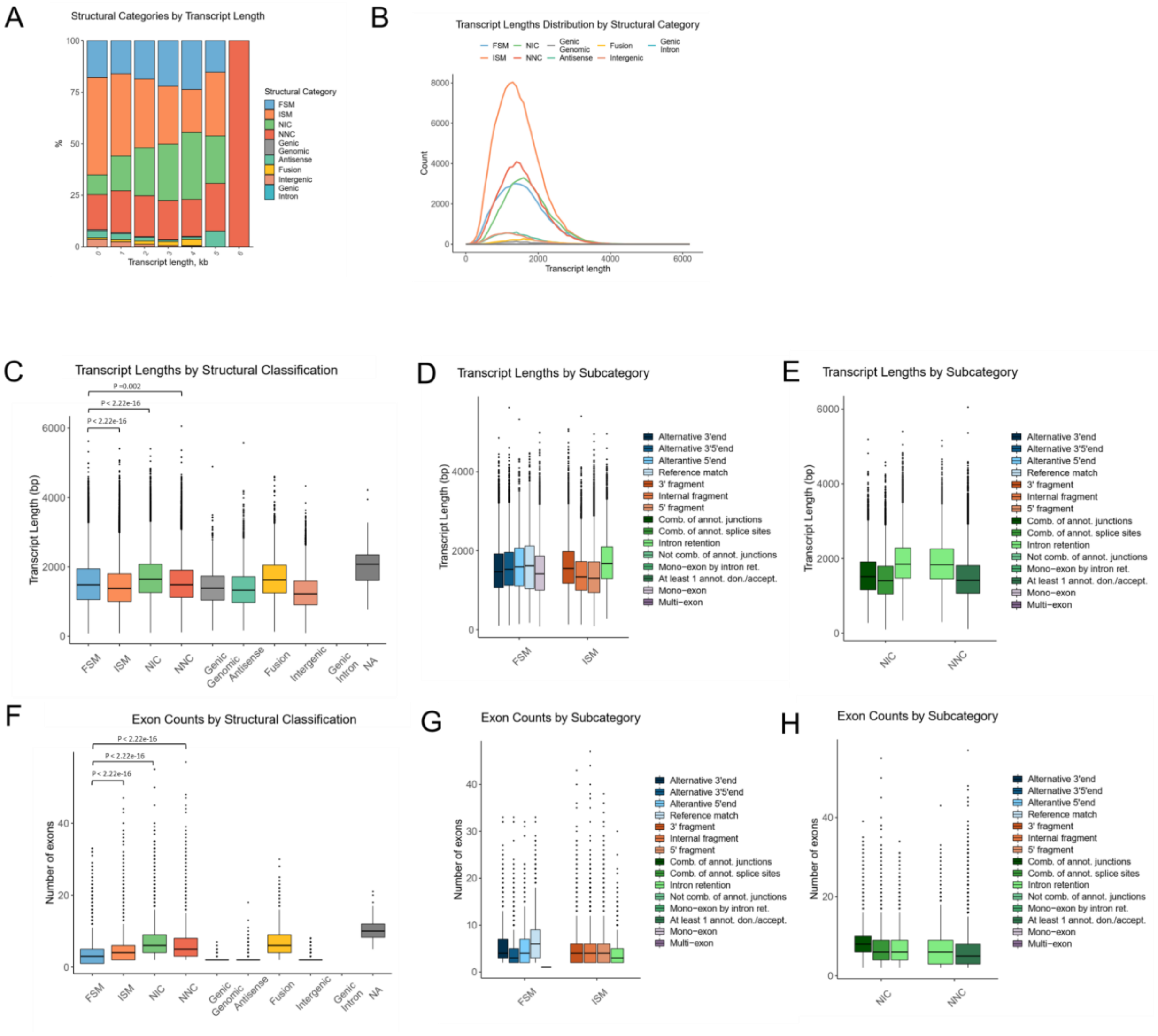


**Supplementary figure 2: Gene and structural isoform characterization.** A) Transcript length grouped by proportion within structural categories. B) Transcript length distribution group by structural categories. Detected transcript length grouped by C) structural categories, D) subcategory of FSM and ISM and E) subcategory of NIC and NNC. Exon counts are grouped by F) structural categories, G) subcategory of FSM and ISM and H) subcategory of NIC and NNC.

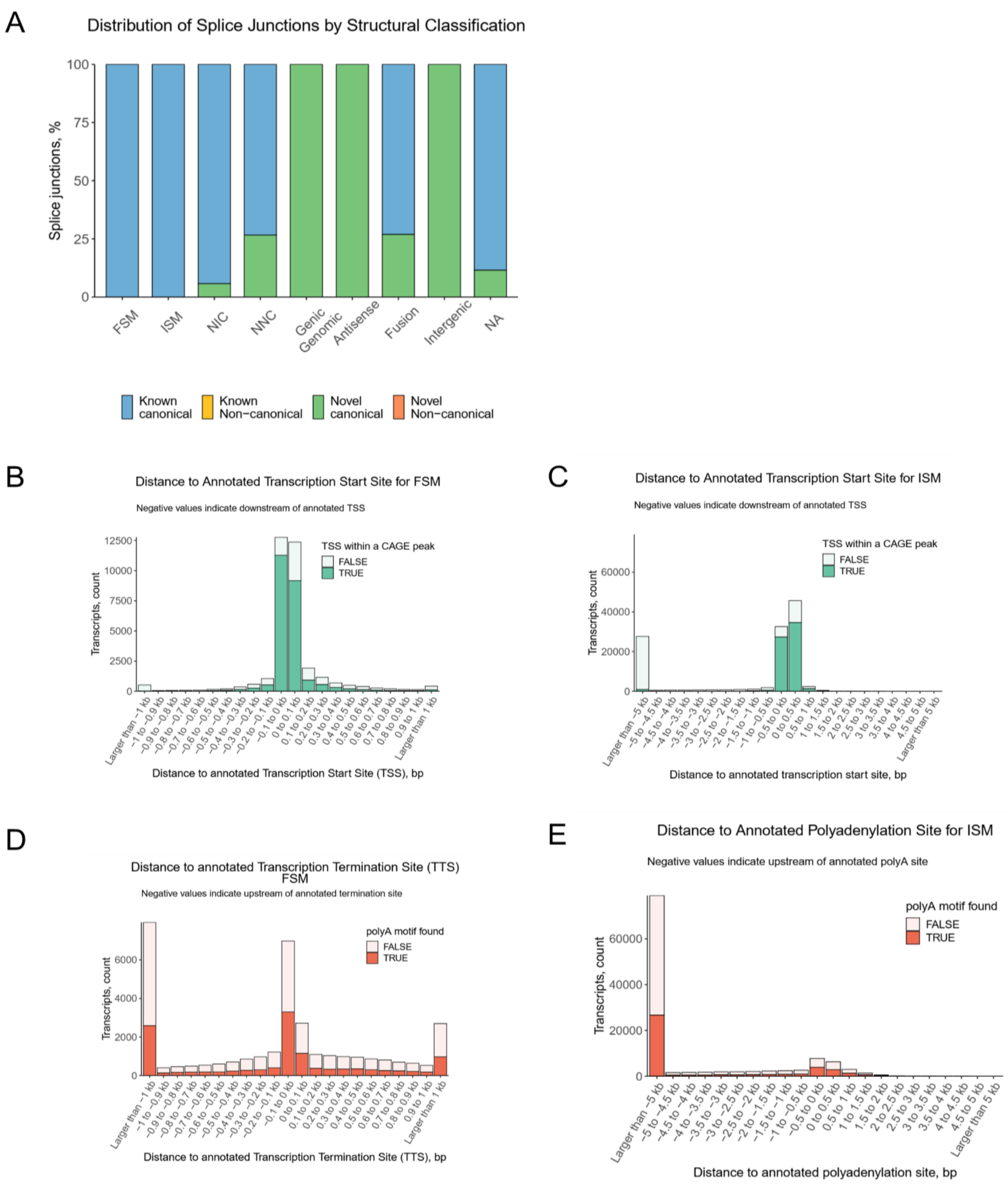


**Supplementary figure 3: Characterization of splice junctions and annotated features of detected long-read transcripts in the human retina.** A) Distribution of splice junctions across transcript categories according to SQANTI classification. Distance to annotated TSS for B) FSM and C) ISM. Distance to annotated transcription termination site (TTS) for D) FSM and E) ISM.

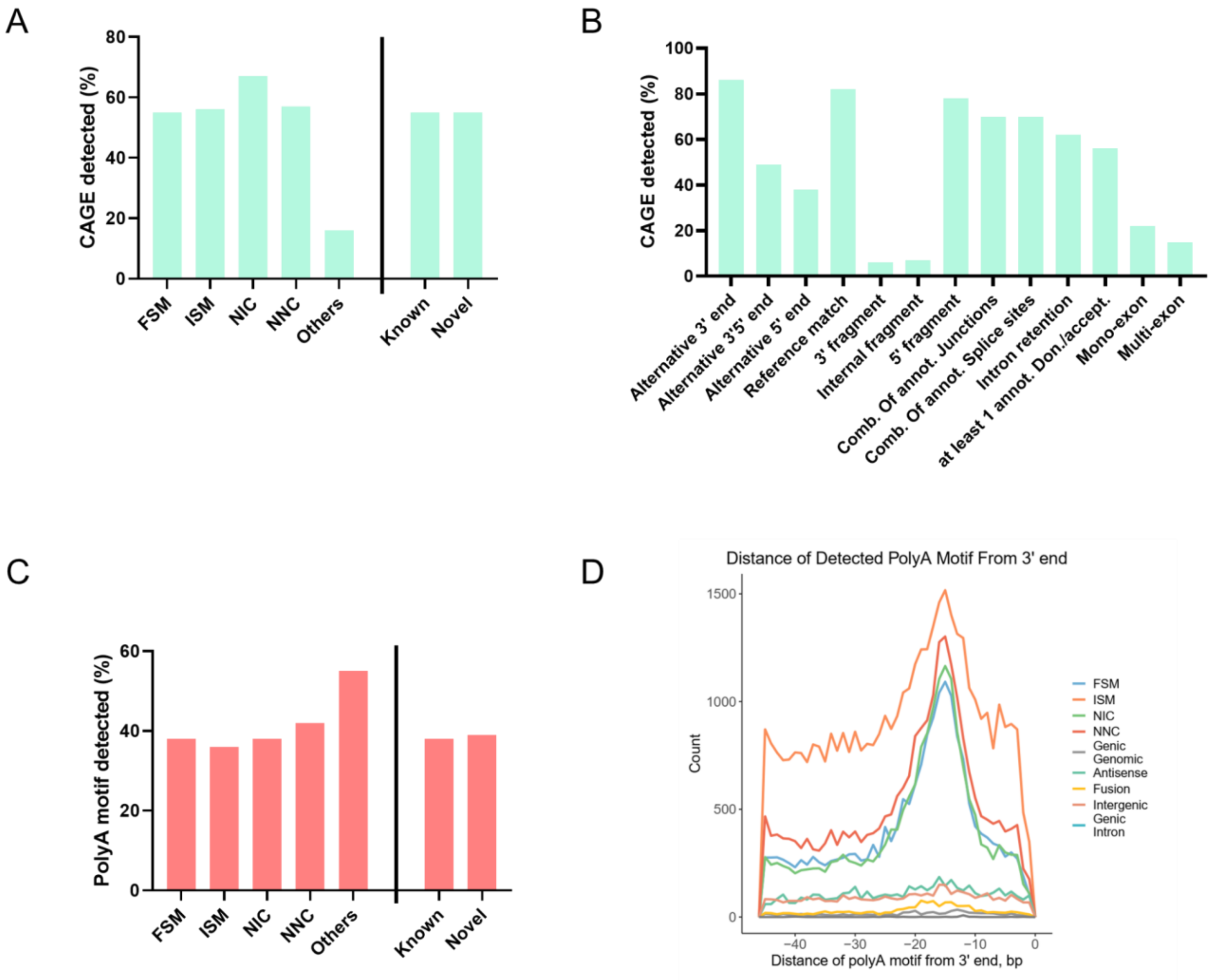


**Supplementary figure 4: Characterization of 5' and 3' features of detected long-read transcripts in the human retina.** Proportion of CAGE detected across transcript categories, including A) known and novel transcripts and B) sub-categories according to SQANTI classification. C) Proportion of poly-A motif detected and D) the distance of poly-A motif from 3' end across transcript categories.

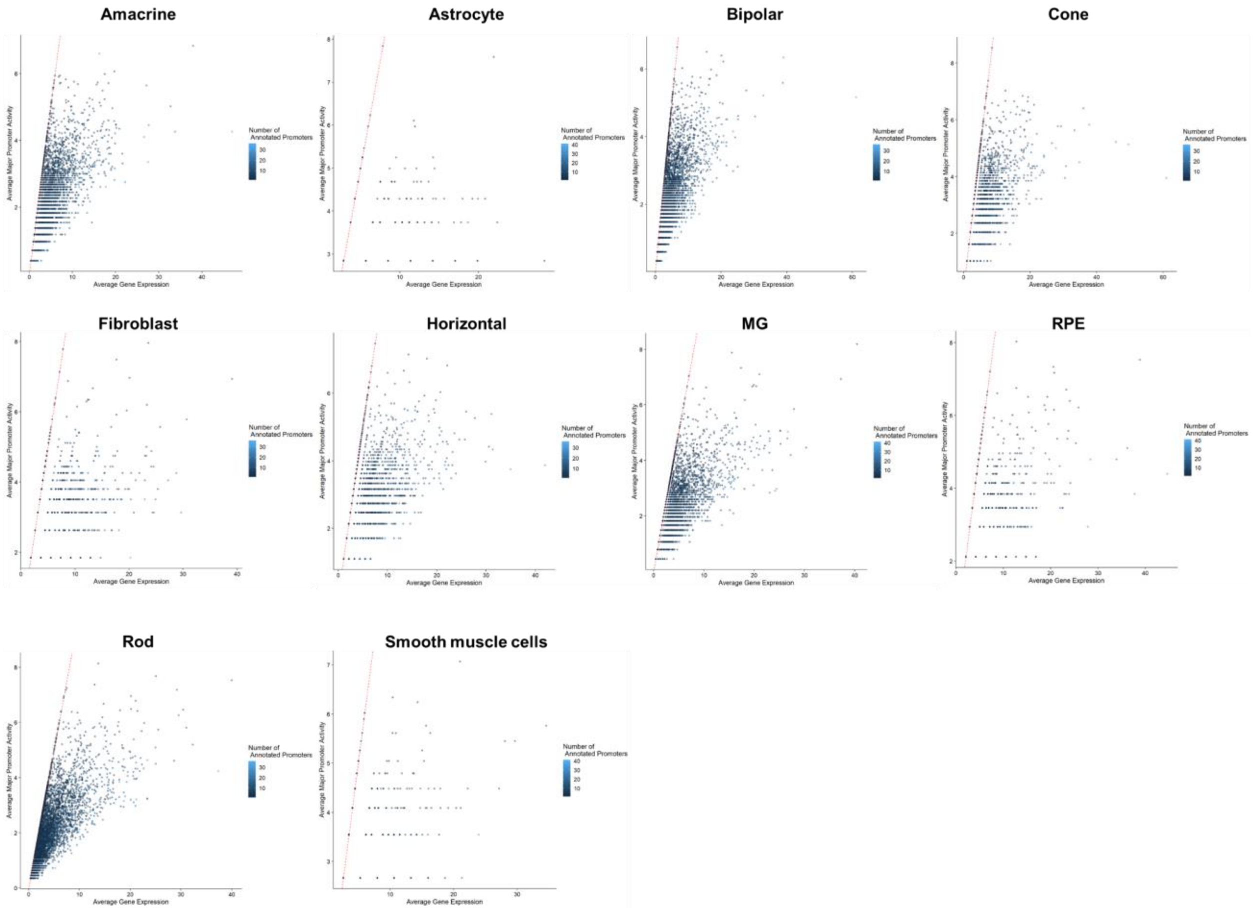


**Supplementary figure 5: Alternative promoter analysis of gene expression in retinal cell types.** Comparison of major promoter activity and gene expression in retinal cell types, calculated by summing over all promoters.

### ABCA4

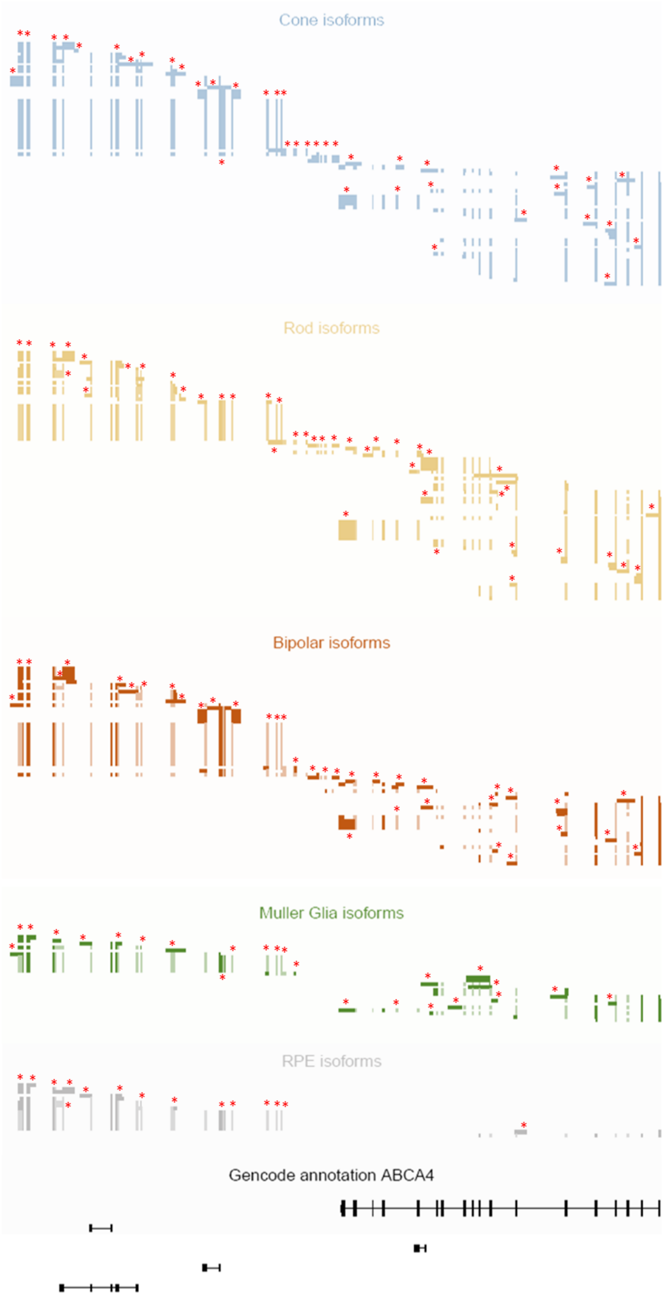


**Supplementary figure 6: Novel exons for *ABCA4* in human retinal cells.** Isoform profiling of *ABCA4* in cones (blue), rods (gold), bipolar cells (brown), Muller glia (green), RPE (grey) compared to annotated isoforms (black). Red asterisks highlighted representative novel exons compared to annotated ones.

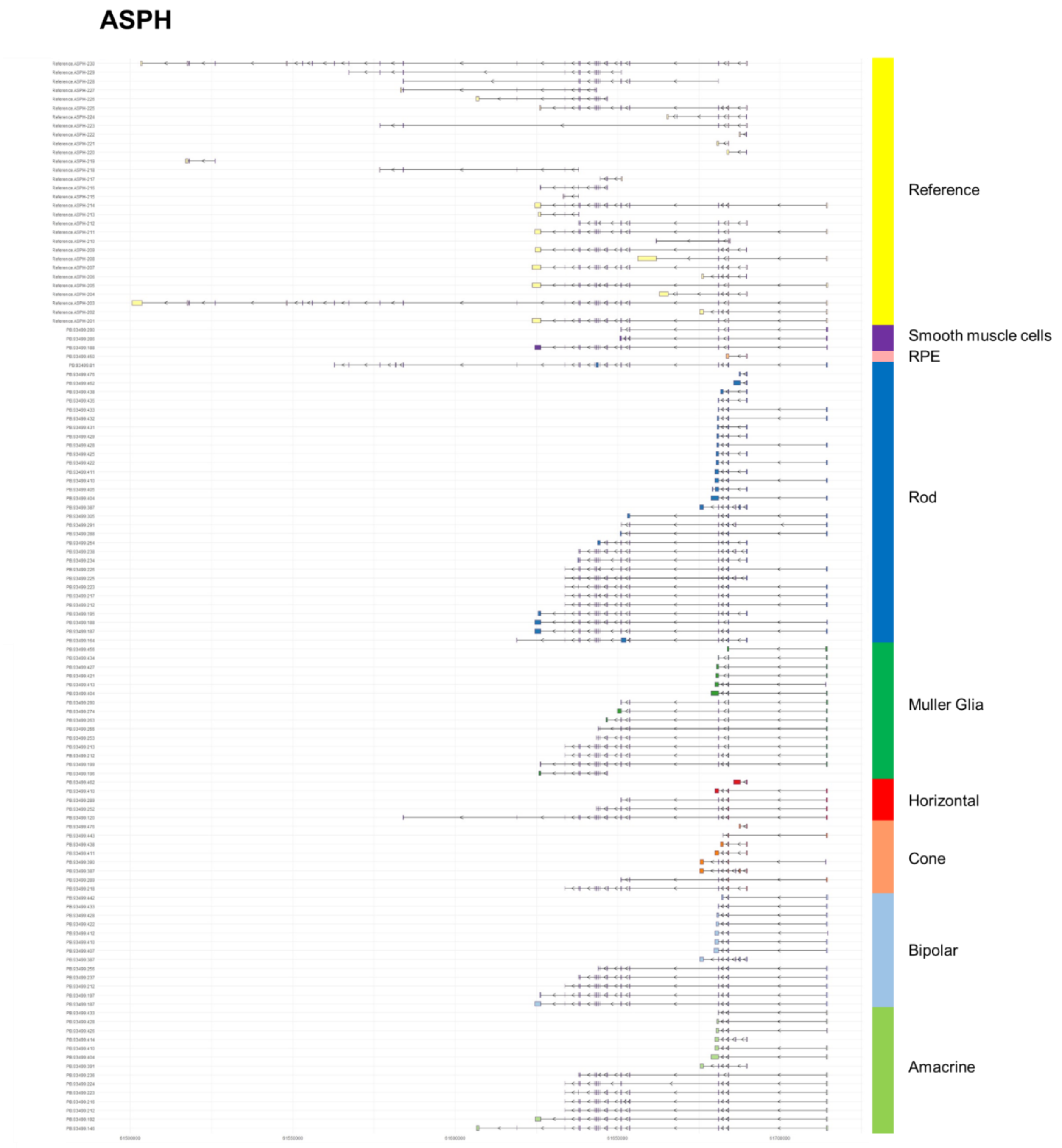


**Supplementary figure 7: *ASPH* isoform profiling in retinal cell types.** Detected full length isoforms of *ASPH* with CAGE peaks and poly-A tails in retinal cell types.

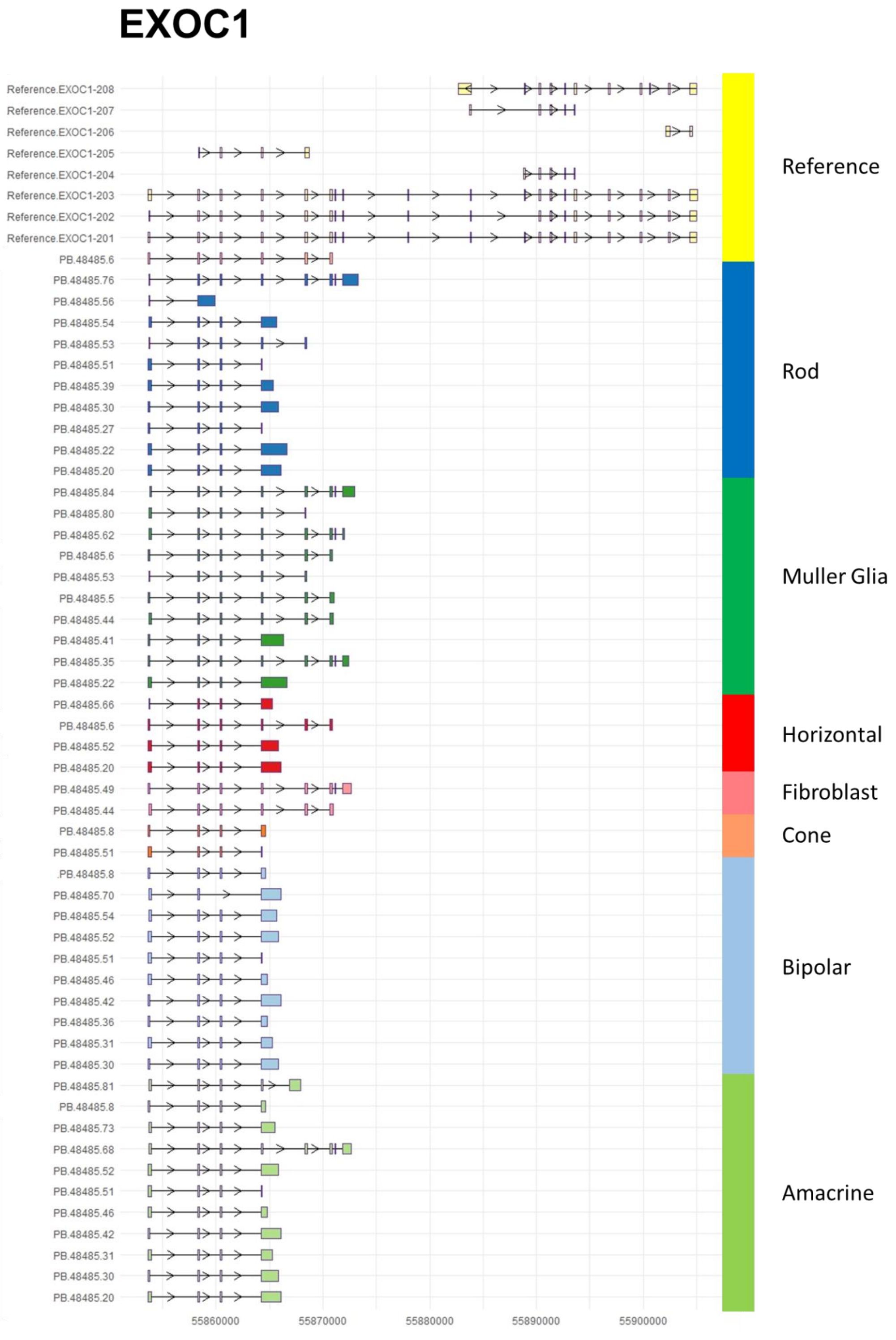


**Supplementary figure 8: *EXOC1* isoform profiling in retinal cell types.** Detected full length isoforms of *EXOC1* with CAGE peaks and poly-A tails in retinal cell types.

**Supplementary table 1:** Sequencing information on retinal samples used in this study

| Sample | Number of nuclei post-QC | Short read sequencing | Output (reads) | Long-read sequencing | Output (CCS reads) | Output (Full-Length Non-Chimeric Reads with Poly-A Tail) |
|---|---|---|---|---|---|---|
| Retina 1 | 8675 | 1x Novaseq SP lane | 424,865,456 | 3x Sequel II SMRT 8M cells | 9,089,764 | 5,094,447 |
| Retina 2 | 8772 | 1x Novaseq SP lane | 468,920,999 | 3x Sequel II SMRT 8M cells | 6,749,605 | 3,779,377 |
| Retina 3 | 7855 | 1x Novaseq SP lane | 500,136,829 | 3x Sequel II SMRT 8M cells | 8,330,198 | 4,752,098 |
